# Control of sensory ectopic spike initiation by descending modulatory projection neurons

**DOI:** 10.1101/025114

**Authors:** Carola Städele, Wolfgang Stein

## Abstract

Descending pathways are important modulators of motor networks and allow the dynamic adjustment of behaviors to changing internal and external conditions. Central pattern generating networks (CPG) have been particularly amenable to study the modulation of motor networks and demonstrated that virtually all levels of information processing are controlled by descending projections. CPGs receive sensory feedback and while it is known that sensory activity can be gated by central pathways, we here present for the first time that descending projection neurons modulate action potential initiation in sensory neurons.

We used the fact that the anterior gastric receptor neuron (AGR), a single-cell bipolar muscle tendon organ in the crustacean stomatogastric nervous system, generates spontaneous ectopic action potentials in its axon. We found that axonal spike initiation is under direct neuromodulatory control by a pair of descending projection neurons. These IV (inferior ventricular) neurons descend from the brain and are known to control CPGs in the stomatogastric ganglion (STG). Activation of the IV neurons elicited a long-lasting decrease in AGR ectopic spike activity. This decrease was only observed when spikes were generated ectopically in the central portion of the axon, i.e. the modulation was specific to the site of spontaneous spike initiation. The decrease could be mimicked by focal application of the IV neuron co-transmitter histamine and IV neuron actions were diminished after blocking H_2_ receptors, indicating a direct descending modulation of the axonal spike initiation site. In contrast, the propagation dynamics of en-passant action potentials were not affected. Descending modulatory projection neurons therefore control axonal spike initiation in sensory neurons without affecting afferent spike propagation to increase the physiological activity repertoire of sensory pathways.

## Introduction

Many rhythmic behaviors such as breathing, swallowing and walking are mediated by central pattern generators (CPGs). CPGs are neural networks that generate rhythmic motor output, and they often do so even in the absence of afferent input (Delcomyn, 1980; Marder and Calabrese, 1996; Nusbaum and Beenhakker, 2002; Grillner, 2009; Selverston, 2010; Harris-Warrick, 2011). Nevertheless, CPGs receive sensory input to adjust their output to changes in the periphery (reviewed in Bässler, 1986; Wolf, 1995; Ausborn et al., 2007; Büschges et al., 2011; Stein, 2014). When this occurs, the stimulus properties are not the only factors to contribute to motor activity. Instead, peripheral and central influences interact to produce the output, making the state of the system and ongoing activity important contributors to stimulus-induced changes in motor output.

A number of mechanisms have been identified that affect sensory pathways, including activity- or state-dependent reduction of afferent spike amplitude (Clarac and Cattaert, 1996; Schmitz and Stein, 2000; Margrie et al., 2001; Barriere et al., 2008), spike conduction block (Burrows and Matheson, 1994; Xiong and Chen, 2002; Lee et al., 2012) and regulation of spike initiation (Evans et al., 2003; Cropper et al., 2004). In addition, encoding of sensory information has been shown to be subject to neuromodulation in several systems (Katz and Frost, 1996; Birmingham, 2001; Birmingham et al., 2003; Mitchell and Johnson, 2003; Dickinson, 2006; Stein, 2009; Nadim and Bucher, 2014), although in most cases the source of the modulation is unknown.

Studies of activity-dependent regulation and modulation are technically difficult and best performed in experimentally advantageous invertebrate preparations with access to individually identified sensory and central pathways. We use one such preparation to study central modulation of the sensory neuron AGR (anterior gastric receptor neuron) which is part of the gastric mill CPG for chewing in the crustacean stomatogastric nervous system (STNS). AGR is a single-cell bipolar muscle-tendon organ (Combes et al., 1995; Smarandache and Stein, 2007). Its soma is located in the stomatogastric ganglion (STG) and protrudes a central axon to upstream modulatory projection neurons in the commissural ganglia (CoG) and a peripheral axon to gastric mill muscles (Fig. 1A, Combes et al., 1995). Part of the normal activity repertoire of AGR is the generation of spontaneous low-frequency tonic ectopic spike activity in its central axon (Smarandache et al., 2008; Daur et al., 2009; Städele and Stein, 2015). Ectopic spiking occurs as soon as peripheral spiking ceases, i.e. at rest and in-between peripherally generated bursts (*in vivo* and *in vitro,* Smarandache et al., 2008; Daur et al., 2009). Those ectopic spikes are propagated orthodromically (towards CoG projection neurons) and antidromically (towards the periphery) (Daur et al., 2009; Städele and Stein, 2015). Ectopic spiking has been identified in sensory and central neurons in a number of different systems and can affect spike initiation (via antidromic propagation) and transmitter release (via orthodromic propagation) (Lena et al., 1993; Pinault, 1995; Cattaert et al., 1999; Bucher and Goaillard, 2011). While the effects of antidromic propagating ectopic spikes in AGR are unknown, Daur et al. (2009) have shown that even small changes in ectopic spike frequency causes pronounced changes in the postsynaptic circuits. To what extent ectopic spike activity is regulated and under neural control is unclear, as well as whether it is influenced by central or sensory pathways.

**Figure 1.**
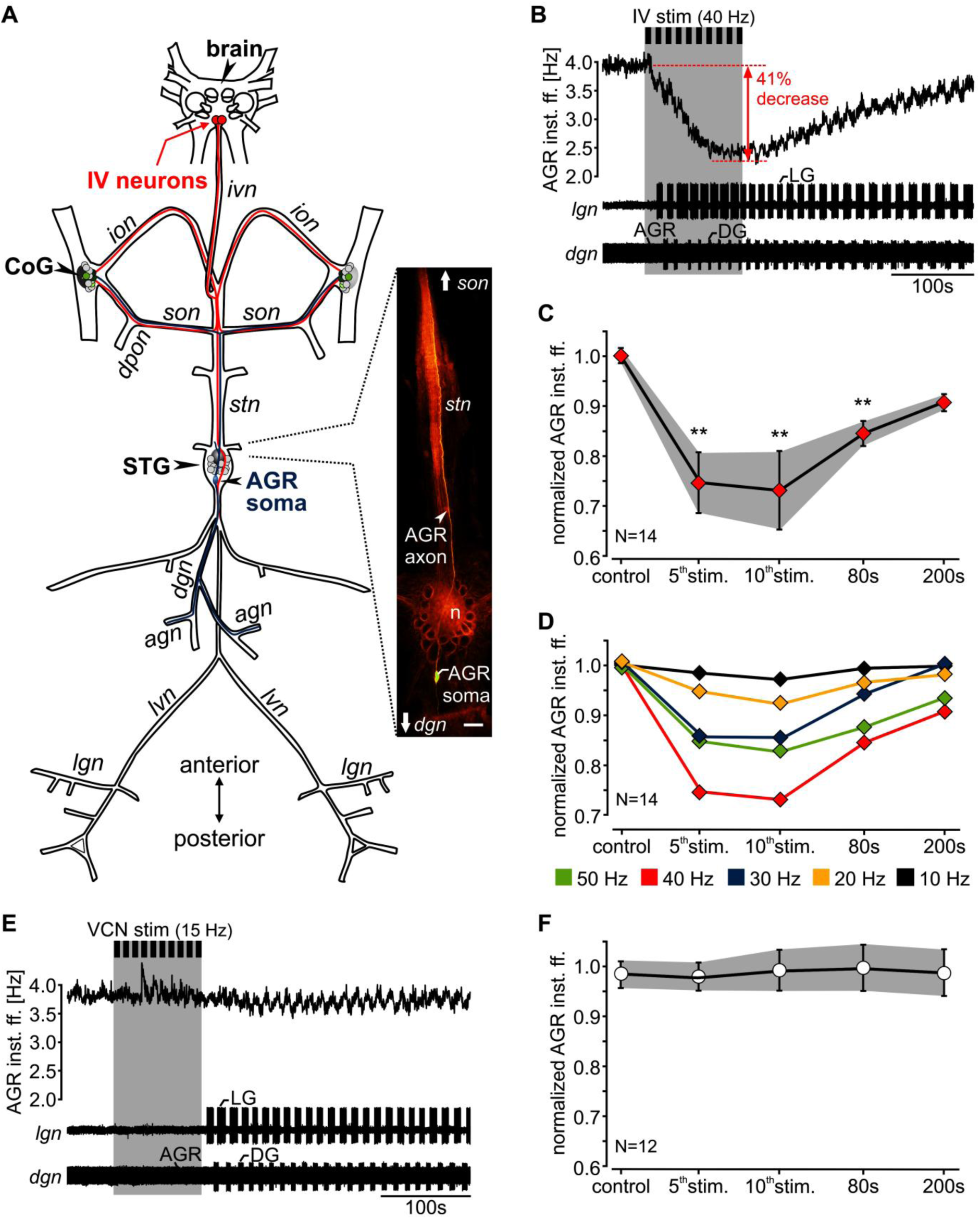
AGR firing frequency diminished during IV neuron stimulation but was not influenced by VCN neurons. (**A**) Schematic of the stomatogastric nervous system. Axonal projections of the paired IV neurons are depicted in red. AGR and its axonal projections are depicted in blue. AGR, the sensory neuron used in this study, is a bipolar neuron with its soma in the STG (see inset on the left) and axonal projections in the *stn* and *dgn*. AGR was visualized via intracellular injection of Alexa Fluor 568. Neural structures were visualized via bath-application of the voltage-sensitive dye Di-4 ANNEPS (Städele et al., 2012). Scale bar, 100 μm. n=neuropil. Nerve names are italicized. Green circles in the CoG represent descending projection neurons. Nerves: *ivn*: inferior ventricular nerve, *ion*: inferior oesophageal nerve, *son*: superior oesophageal nerve, *dpon*: dorsal posterior oesophageal nerve, *stn*: stomatogastric nerve, *dgn*: dorsal gastric nerve; *agn*: anterior gastric nerve, *lvn*: lateral ventricular nerve, *lgn*: lateral gastric nerve. Ganglia: STG: stomatogastric ganglion, CoG: commissural ganglion, brain: supraesophageal ganglion. Neurons: AGR: anterior gastric receptor neuron, IV: inferior ventricular neurons. Adapted from Hedrich and Stein (2008). (**B**) AGR instantaneous firing frequency (inst. ff.) before and during IV neuron stimulation (grey bar). The extracellular nerve recordings of the *lgn* and *dgn* show the response of the STG gastric mill CPG to IV stimulation. Consecutive stimulation of the IV neurons with 40 Hz elicited a gastric mill rhythm (note the alternating activity of LG on the *lgn* and DG on the *dgn)* and a concurrent 41% decrease in AGR inst. ff. (**C**) Average time course of the change in AGR firing frequency during IV neuron stimulation (10 consecutive trains, 40 Hz stimulation frequency) for 14 experiments. AGR firing frequency was normalized to the frequency measured 100 sec before IV neuron stimulation (=baseline). Control refers to the frequency measured immediately before the stimulation. Shown are mean values ± SD (grey area). AGR inst. ff. was significantly diminished for all measurements after the 5^th^ IV neuron stimulation train and up to 80 sec after the end of the stimulation. One way RM ANOVA, F(4,55)=49.21, p<0.001, Holm-Sidak post-hoc test with p<0.01 significance level, N=14. (**D**) Average time course of the change in normalized AGR inst. ff. during IV neuron stimulation with 10 to 50 Hz. Shown are the mean values for 14 experiments. (**E**) AGR inst. ff. before and during VCN stimulation (grey bar) for the same experiment as shown in A. VCN stimulation did not affect AGR inst. ff., but started a gastric mill rhythm (see LG on the *lgn* and DG on the *dgn*). (**F**) Average time course of normalized AGR inst. ff. in response to VCN stimulation. Shown are means ± SD for 12 preparations. VCN stimulation did not cause a significant change in AGR firing frequency (one way RM ANOVA, F(4,47)=0.65, p=0.66, N=12).

We have previously shown that ectopic firing in AGR appears to be weakly excited by gastric mill motor neurons in the STG (Goldsmith et al., 2014; Städele and Stein, 2015). Here, we show that axonal spike initiation in AGR is directly modified by descending modulatory projection neurons that control the CPGs in the STG. We found that the inferior ventricular neurons (IV, Christie et al., 2004; Hedrich and Stein, 2008), a pair of descending chemosensory projection neurons located in the supraesophageal ganglion (brain, Fig. 1A), elicited a long-lasting decrease in AGR spike frequency. This decrease could only be observed when spikes were generated ectopically in the central axon (close to the STG neuropil). The IV neuron effect was mediated via release of Histamine at the central spike initiation zone. We thus demonstrate that the direct control of ectopic spiking in a sensory neuron can increase the physiological repertoire of sensory feedback without the need for changing circuit connectivity or synaptic weighting.

## Materials And Methods

### Dissection

Adult crabs (*Cancer borealis*) were purchased from The Fresh Lobster Company (Gloucester, MA) or Ocean Resources Inc. (Sedgwick, ME) and kept in tanks with artificial sea water (salt content ∼1.025g/cm^3^, Instant Ocean Sea Salt Mix, Blacksburg, VA) at 11°C and a 12-hour light-dark cycle. Before dissection, animals were anesthetized on ice for 20-40 minutes. All experiments were performed *in vitro* on isolated nervous systems. The stomatogastric nervous system (STNS) including the supraesophageal ganglion (brain, Fig. 1A) was isolated from the animal following standard procedures, pinned out in a silicone lined (Wacker) petri dish and continuously superfused (7-12 ml/min) with physiological saline (10-11°C). Experiments were performed on fully intact and decentralized preparations. In the latter, the CoGs were removed by transecting the paired *ion* and *son*.

All animal procedures were performed in accordance with the Illinois State University animal care committee’s regulations. We adhered to general animal welfare considerations regarding humane care and use of animals and Guidelines laid down by the NIH. Animals were sacrificed on ice, a method recognized as acceptable under the AVMA guidelines for euthanasia of invertebrates.

### Solutions and modulators

*C. borealis* saline was composed of (in mM) 440 NaCl, 26 MgCl_2_, 13 CaCl_2_, 11 KCl, 11.2 Trisma base, 5 Maleic acid, pH 7.4-7.6 (Sigma Aldrich). In some experiments, low Ca^2+^ saline was applied to the posterior part of the *stn* (close to the STG neuropil) to block chemical transmission. Low Ca^2+^ saline was composed of 440 NaCl, 11 KCl, 26 MgCl_2,_ 0.1 CaCl_2,_ 11.2 Trisma base, 5.1 Maleic acid, 12.9 MnCl_2_, pH 7.4-7.5. High-divalent saline (“HiDi”) was composed of 439 NaCl, 130 MgCl_2_, 64.5 CaCl_2_, 11 KCl, 12.4 Trisma base, 5 Maleic acid and contained five times the amount of Ca^2+^ and Mg^2+^ than regular saline. HiDi was superfused to the posterior half of the *stn* including the STG neuropil. In these experiments, AGR spike activity was monitored extracellularly in the anterior part of the *stn* (close to the *stn/son* junction). which was not affected by HiDi.

### Extracellular and intracellular recordings

Previously, Goldsmith et al. (2014) demonstrated that removing the sheath of the stomatogastric ganglion (STG) influences modulation of AGRs ectopic spiking. Thus, if not stated otherwise, all experiments were carried out using non-desheathed nervous system preparations and extracellular recording techniques. For extracellular recordings, petroleum jelly-wells were built to electrically isolate a small part of the nerve from the surrounding bath. One of two stainless steel electrodes was placed inside the well to record neuronal activity of all axons projecting through a particular nerve. The other wire was placed in the bath as reference electrode. Extracellular signals were recorded, filtered and amplified through AM Systems amplifier (Model 1700, Carlsborg, WA). Files were recorded, saved and analyzed using Spike2 Software (version 7.11; CED, UK) at 10 kHz. The activity of AGR was monitored on multiple extracellular recordings simultaneously, namely on the *stn*, *dgn*, and the *son* (see Fig. 1A). AGR activity was measured as instantaneous firing frequency (inst. ff.) as determined by the reciprocal of the interspike interval.

### Extracellular axon stimulation

Retrograde extracellular nerve stimulations were performed using a Master-8 pulse stimulator (A.M.P.I., Israel) controlled by self-programmed Spike2 scripts. Extracellular activation of sensory modalities is well established in the STNS and we used standard stimulation protocols to activate the IV neurons (Hedrich and Stein, 2008), the ventral cardiac neurons (VCN, Beenhakker et al., 2004) and AGR (Smarandache and Stein, 2007). A petroleum jelly well was built around a nerve containing the axons of the neuron of interest. One of two stainless steel stimulation electrodes was placed inside the compartment, the other was placed outside. The IV neurons were activated via extracellular stimulation of the inferior ventricular nerve (*ivn*, see Fig. 1A) with 10 consecutive stimulus trains, 10 to 50 Hz stimulation frequency, 6 sec stimulus trains, 6 sec intertrain intervals, 1 ms pulse duration, 0.5 to 2 Volt stimulation voltage (Hedrich et al., 2009; Hedrich et al., 2011). The VCNs, were activated via extracellular stimulation of the paired dorsal posterior oesophageal nerves (*dpon*) with 10 consecutive stimulus trains, 15 Hz stimulation frequency, 6 sec stimulus trains, 4 sec intertrain intervals, 1 ms pulse duration, 2 to 3 Volt stimulation voltage (Beenhakker et al., 2004). In all experiments both *dpons* were stimulated simultaneously using different channels on the Master-8 stimulator. AGR was stimulated on the anterior gastric nerve (*agn,* Fig. 1A), a side branch leaving the *dgn* that exclusively contains the AGR axon. To detect differences in spike failures before and during IV neuron modulation, the *agn* was stimulated with 10 consecutive trains, 10 to 40 Hz stimulation frequency, 9 sec stimulus trains, 9 sec intertrain intervals, 1 ms pulse duration and 0.5 to 1 Volt. To determine changes in spike conduction velocity we used 5 consecutive trains of 15 Hz stimulation frequency, each train with 28 pulses, 6 to 9 sec intertrain interval and 1 ms pulse duration.

### Neuromodulator and antagonist application

Neuromodulators and antagonists were diluted in ultrapure water (18.3 MΩ) and stored in small quantities as concentrated stock solutions at -20°C. Immediately before an experiment the neuromodulators were diluted in saline to the desired concentration. Concentrations varied between neuromodulators and are stated in the text. Histamine dihydrochloride (Sigma Aldrich), FMRF-like peptide F1 (TNRNFLRF-NH_2_, Bachem) and cimetidine hydrochloride (Sigma Aldrich) were bought commercially. Cimetidine hydrochloride was dissolved in dimethyl sulfoxide (DMSO) and protected from light throughout the length of the experiment. The two Orcokinin isoforms used ([Ala^13^] and [Val^13^] orcokinin, Li et al., 2002) were a gift from Lingjun Li (University of Wisconsin at Madison, WI, USA). Neuromodulators and antagonists were applied selectively to the posterior part of the *stn* where AGR’s ectopic spike initiation zone is located (Städele and Stein, 2015). A petroleum-jelly well was used to isolate the application site from the rest of the nervous system. The well had an inner diameter of approximately 300-400 μm. Neuromodulators and antagonists were cooled to 10- 12°C and manually applied into the well using a 1 ml syringe with an injection needle. To exclude temperature-induced changes in AGR frequency, saline with the same temperature as the neuromodulators/antagonists was applied 5 minutes before each application. Measurements were taken in steady-state (5 minutes after neuromodulator/antagonist wash in). Modulators were washed out for 20 to 40 minutes with continuous superfusion of cooled saline.

Similar to observations by Goldsmith et al. (2014) we found that removing the connective tissue from the STG caused subtle changes in AGR activity, namely a diminishment of AGR’s response to IV neuron stimulation. Thus, all neuromodulators and antagonists were applied to non-desheathed preparations. To test whether a particular substance can penetrate the connective tissue, we performed control experiments where we applied the modulator/antagonist to the non-desheathed STG and observed whether or not the substance affected the STG circuits in a similar fashion as previously published for desheathed ganglia (FMRF-like peptide F1: Christie et al., 2004, Histamine: Stein et al., 2007, Orcokinin: Li et al., 2002). For each substance, several concentrations were tested to determine the effective concentration.

### Data Analysis, statistics and figure making

Data were analyzed using scripts for Spike2 (available on www.neurobiologie.de/spike2) and by using built-in software functions. For spike detection a voltage threshold was used and the subsequent maximum or minimum voltage deflection of the signal passing through this threshold was used as trigger. In cases where extracellular stimulus artifacts or the action potentials of other neurons obscured the neuron of interest, obscuring signals were eliminated from recordings by subtracting the average stimulus artifact with Spike2. For spreadsheet analysis, Excel (version 2010 for Windows, Microsoft) or SigmaPlot (version 12 for Windows, Systat Software GmbH, Erkrath, Germany) were used. Normally distributed data are given as mean ± SD. “N” denotes the number of animals, while “n” is the number of trials. Significant differences are indicated using * (p<0.05), ** (p<0.01), *** (p<0.001). Detailed information about statistical tests is given in the corresponding figure legend. Final figures were prepared with CorelDRAW Graphics Suite (version X7, Corel Corporation, Ottawa, ON, Canada).

## Results

We have previously shown that ectopic spikes in AGR are initiated in posterior parts of the *stn,* in vicinity to the anterior border of the STG. Those ectopic spikes are weakly excited by the STG gastric mill (GM) motor neurons during ongoing gastric mill rhythms (Städele and Stein, 2015). The STG is innervated by a set of descending modulatory projection neurons from the commissural and oesophageal ganglia as well as from the brain. These projections initiate or modulate the pyloric and gastric mill CPGs (summarized in Marder and Bucher, 2001; Selverston et al., 2009; Stein, 2009; Blitz and Nusbaum, 2011). The AGR axon projects through the STG (Fig. 1A) and albeit no connections from AGR to descending projection neurons have been identified, Goeritz et al. (2013) suggested putative chemical connections to an unknown modulatory projection neuron.

### AGR’s ectopic spike activity is influenced by descending projection neurons

To test whether ectopic spike initiation in AGR is under neural control by descending projections, we selectively activated the two chemosensory IV neurons. This pair of neurons descends from the brain and innervates both the STG and the commissural ganglia (Fig. 1A). To activate the IV neurons we performed repetitive extracellular nerve stimulations (according to their *in vivo* firing pattern, Hedrich and Stein, 2008) of the inferior ventricular nerve (*ivn*) through which the IV neuron axons project (Fig. 1A). We monitored AGR ectopic spike frequency extracellularly on several nerves (the peripheral dorsal gastric nerve (*dgn*), the central stomatogastric nerve (*stn*) and close to CoG projection neurons on the superior oesophageal nerve (*son*)). Figure 1B shows the response of AGR and the gastric mill motor circuit to IV neuron stimulation. IV neurons were stimulated with 40 Hz and 10 consecutive trains of 6 sec train/intertrain duration. IV neuron stimulation induced a strong decrease in AGR instantaneous firing frequency (from 3.9 Hz to 2.3 Hz, Δf=1.6 Hz) that outlasted the stimulation for more than 300 seconds. Across preparations, IV neuron stimulation caused a significantly decrease in AGR firing frequency by 26.7±7.9% (Fig. 1C, one way RM ANOVA, p<0.01, N=14).

Additionally, as described previously (Hedrich and Stein, 2008) IV neuron stimulation started a long-lasting gastric mill rhythm with alternating burst activity of the lateral gastric (LG) neuron on the lateral gastric nerve (*lgn*) and the dorsal gastric (DG) neuron on the *dgn* (Fig. 1B, bottom). The gastric mill rhythm was accompanied by small rhythmic frequency changes in AGR which are likely to be mediated by the gastric mill (GM) motor neurons (Goldsmith et al., 2014; Städele and Stein, 2015). We found that the decrease in AGR firing frequency could be prolonged when the IV neurons were stimulated for longer durations (up to 40 stimulus trains, N=5, data not shown). In these cases, the beginning of the recovery back to baseline frequency was also delayed until the end of the stimulation.

To characterize the time course of the IV neuron-mediated effect we measured AGR firing frequency before IV stimulation (control), during the 5^th^ and 10^th^ stimulation train and 80 sec and 200 sec after the last stimulation (Fig. 1C). Across animals (N=14) we found that the IV neuron-mediated effect had a slow time course: AGR firing frequency was significantly diminished starting with the 5^th^ stimulation and up to 80 sec after the last IV stimulation (p<0.01, one way RM ANOVA, N=14). On average, AGR firing frequency was most diminished at the end of the IV stimulation (= 10^th^ stim). The response of AGR to IV neuron stimulation depended on stimulation frequency. When we stimulated the IV neurons with frequencies between 10 to 50 Hz (Fig. 1D, 10 Hz step intervals) stimulation elicited gastric mill rhythms at all frequencies, with higher stimulation frequencies causing longer lasting and stronger rhythms. However, although 50 Hz stimulation elicited the strongest gastric mill rhythms, 40 Hz elicited the strongest reduction in AGR firing frequency. Higher and lower frequencies were less effective. Significant responses could be observed at 20 Hz, but lower frequencies (10 Hz) had no effect on AGR (AGR inst. ff. after 5^th^ stim. with 20 Hz was significant different from control, one way RM ANOVA, F(4,55)=3.73, p<0.01, N=14).

### Axonal spike initiation is directly controlled via chemical synaptic transmission

The IV neurons exert their actions on the STG motor circuits directly via direct histaminergic inhibition of STG circuits and indirectly via activation of modulatory projection neurons in the commissural ganglia (Christie et al., 2004; Hedrich et al., 2009). Their latter actions are required for activation of the gastric mill rhythm and involve two identified projection neurons, MCN1 (modulatory commissural neuron 1) and CPN2 (commissural projection neuron 2) (Christie et al., 2004; Hedrich et al., 2009; Hedrich et al., 2011). The fact that the observed decrease in AGR firing frequency during IV neuron stimulation was always accompanied by a gastric mill rhythm suggested that the commissural neurons might be involved. To test whether commissural projections contributed to the response of AGR, we elicited gastric mill rhythms that involved the same two projection neurons (but not IV neurons) via activation of the mechanosensory VCN neurons (Beenhakker et al., 2004).

For that purpose, we activated the VCNs via extracellularly stimulation of the dorsal posterior oesophageal nerve (*dpon*, Fig. 1A) with 10 consecutive stimulus trains, each with 15 Hz stimulation frequency (according to Beenhakker et al., 2004). VCN stimulation started a strong and long-lasting gastric mill rhythm (note the alternating activity of LG and DG; Fig. 1E) which was accompanied by rhythmic firing frequency changes in AGR (similarly as previously reported; Goldsmith et al., 2014; Städele and Stein, 2015). However, there was no consistent or long-lasting decrease in AGR firing frequency. We found this to be true for all preparations tested (Fig. 1F, N=8). With the exception of the small rhythmic firing frequency changes, AGR firing frequency remained unaffected during and after VCN stimulation. This indicated that the decrease in AGR firing frequency was specific to the IV neurons and not dependent on the activation of MCN1 and CPN2.

To further scrutinize this result, we completely removed the CoG projection neurons by transecting the *ions* and *sons.* This left the direct connection of the IV neurons to the STG intact but eliminated indirect effects via CoG neurons. Figure 2A shows the AGR ectopic firing frequency before and after CoG transection. Recordings are from the same preparation. IV neuron activation elicited similar decreases in AGR firing frequency in both conditions. This was consistent across preparations (Fig. 2C) and there was no significant change in AGR firing frequency decrease when the CoGs were transected in comparison to control (p=0.85, one way RM ANOVA, N=8). We also noted that in this condition the smaller AGR frequency oscillations were absent.

**Figure 2.**
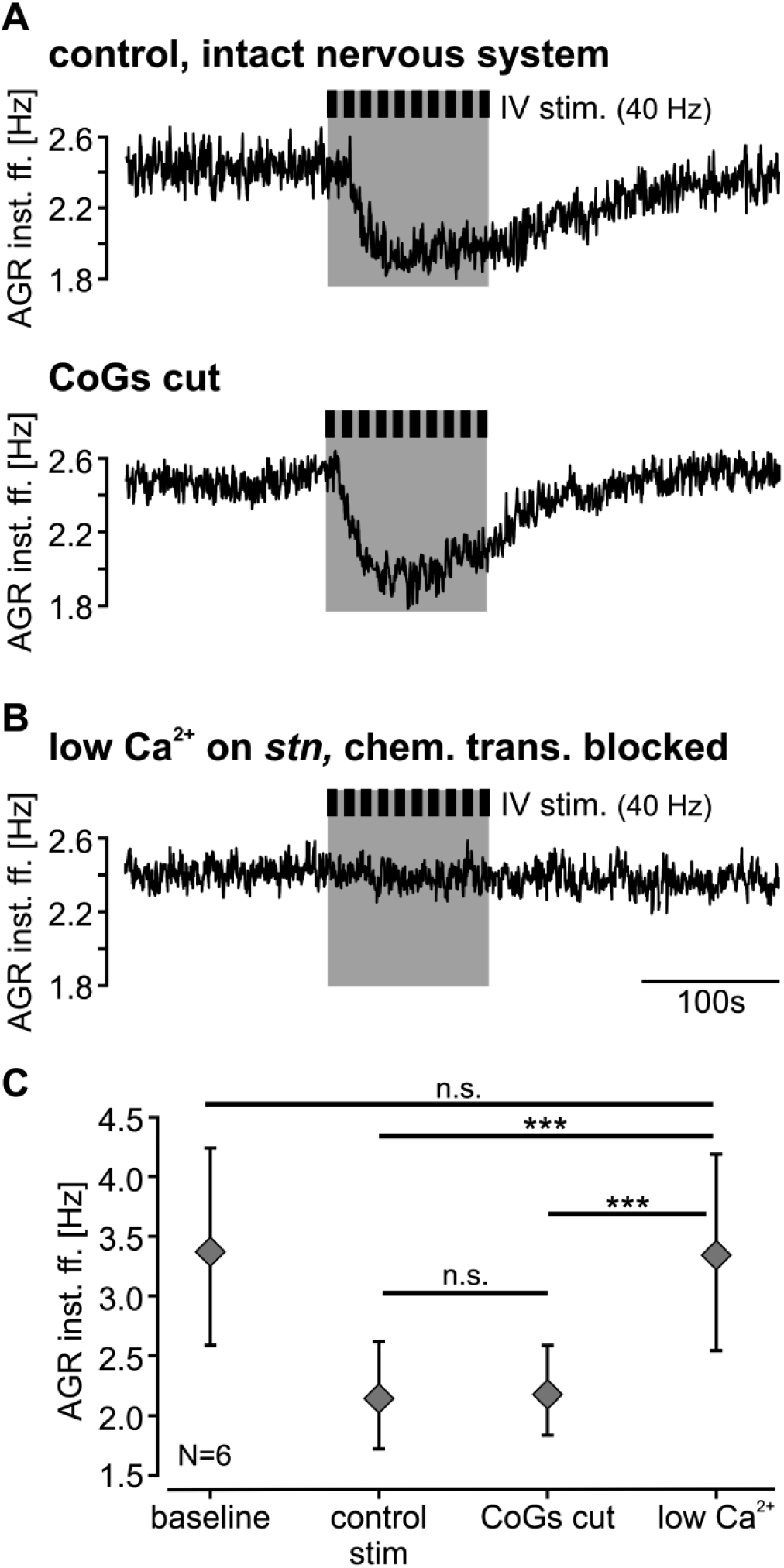
The IV neurons directly diminish ectopic spiking via chemical synaptic transmission. (**A, B**) AGR inst. ff. during IV neuron activation in the intact nervous system (**A**, top) and after CoG transection (**A**, bottom), and after block of chemical transmission via application of low Ca^2+^ saline to the posterior *stn* (**B**). Recordings are from the same preparation. (**C**) Average change in AGR inst. ff. during IV neuron stimulation in control, after CoG transection and after chemical transmission was blocked (low Ca^2+^) for 6 tested preparations. Shown are means ± SD. Baseline refers to the frequency measured before IV stimulation. One way RM ANOVA, F(3,15)=32.36, p<0.001, Holm-Sidak post-hoc test with p<0.01 significance level, N=14). n.s. = not significant different with p>0.8.

The above experiments indicated that the decrease in AGR firing frequency could be mediated via a direct influence of the IV neurons onto AGR. The IV neurons are modulatory projection neurons known to contain at least two co-transmitters, namely Histamine (Stein et al., 2007) and FMRF-like peptide F1 (Christie et al., 2004). Li et al. (2002) suggested that the IV neurons might potentially also contain different Orcokinin isoforms.

To test if the diminishment in AGR firing frequency was mediated by one of the IV cotransmitters, we first examined whether the decrease in AGR firing frequency was chemically transmitted. For this, we blocked chemical transmission at AGR’s ectopic spike initiation zone (eSIZ) by reducing the extracellular calcium concentration via focal application of low Ca^2+^ saline to the posterior part of the *stn.* When chemical transmission was blocked, IV neuron stimulation did no longer cause a decrease in AGR firing frequency (Fig. 2B). We found this to be true for all preparations tested (N=6, Fig. 2C).

We then tested the effects of the IV neuron co-transmitters on AGR by bath applying them individually to the posterior *stn*. We used Histamine (HA), FMRF-like peptide F1 and two different Orcokinin isoforms ([Ala^13^] and [Val^13^] orcokinin, Li et al., 2002) at different concentrations. Figures 3A-C show the AGR response to modulator application. Measurements were taken in steady-state, i.e. 5 minutes after application. The two Orcokinin isoforms never elicited a change in AGR firing frequency (Figs. 3A and D, N=5) despite the fact that both isoforms influence the pyloric and gastric mill CPGs when applied to the STG (Li et al., 2002). This was true for all concentrations tested (1 to 100 μM, N=5). Application of 100 μM FMRF-like peptide F1 to the posterior part of the *stn* excited AGR and elicited a strong increase in firing frequency (Fig. 3B). This increase was concentration-dependent. 1 μM FMRF-like peptide already caused an increase in AGR firing frequency, but the effect was much smaller. On average we found that 100 μM FMRF-like peptide caused a significant increase in AGR firing frequency by 32±15% (Fig. 3D middle, from 2.76±0.90 Hz to 3.58 ±1.02 Hz, p<0.001, paired t-test, N=6). In all cases, the time constant of the increase was slow and steady state was reached after about 120 sec. Application of histamine (10mM), in contrast, caused a strong diminishment in AGR firing frequency (Fig. 3C) with a similar time course as seen during IV neuron activation (for comparison see Fig. 1B). On average, histamine caused a significant decrease in AGR firing frequency by 28±9% (Fig. 3D bottom, from 3.08±0.8 Hz to 2.43±0.91 Hz, p<0.001, paired t-test, N=13) that outlasted the application by about 250 seconds.

**Figure 3.**
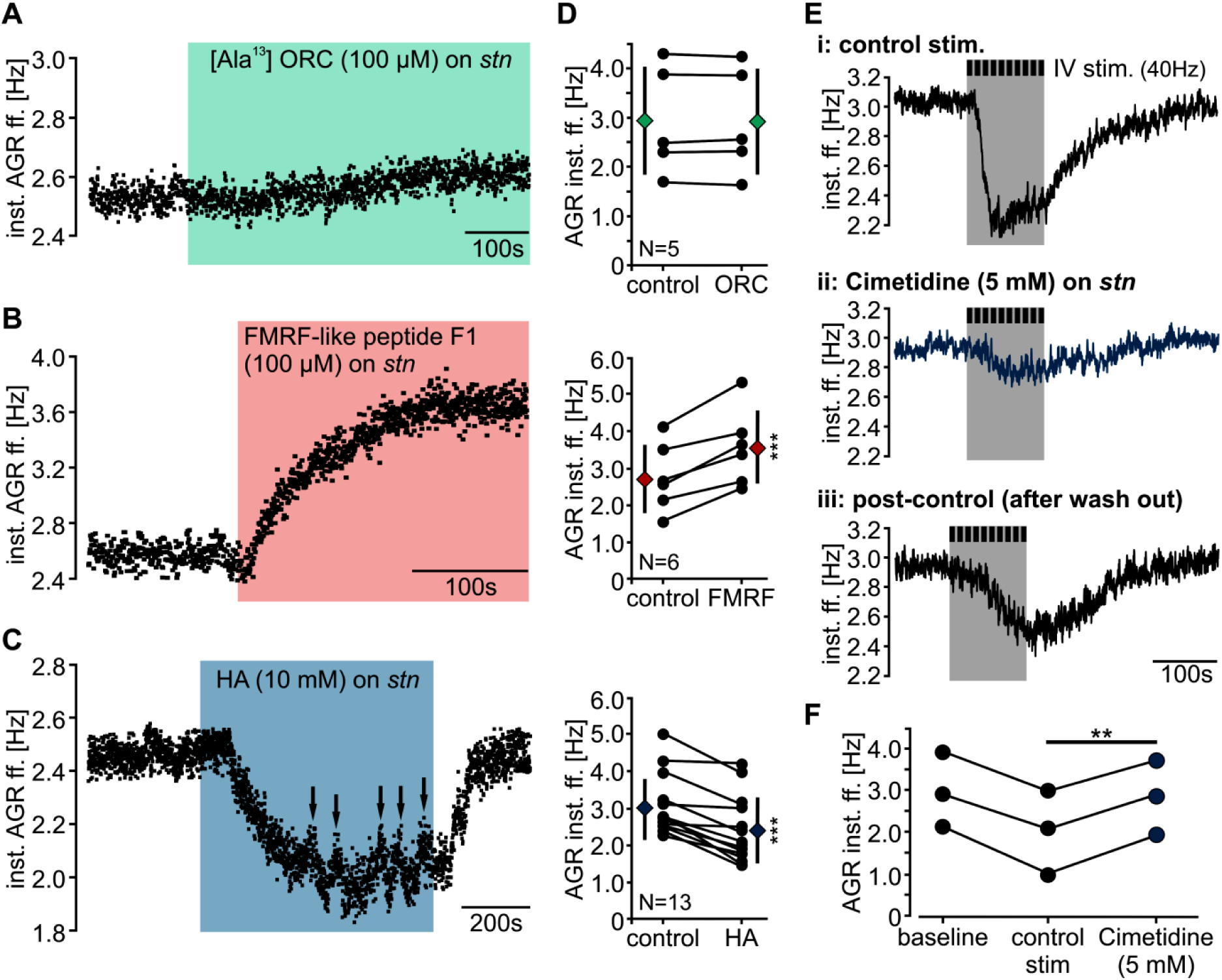
The IV neuron co-transmitter histamine diminishes the inst. AGR. ff. mainly *via* acting on H_*2*_ receptors. (**A-C**) AGR inst. ff. in response to [Ala^13^] orcokinin application (**A**, green bar), FMRF-like peptide F1 (**B**) and histamine (**C**) to the posterior *stn*. Recordings are from different preparations and are scaled differently. The arrows in C indicate a switch in the site of spike initiation. (**D**) Average change in AGR inst. ff. before and during application of the different IV neuron co-transmitters to the posterior *stn*. Black circles represent individual data, diamonds represent means ± SD. AGR. inst. ff. changed significantly (paired t-test, p<0.001) after application of FMRF-like peptide F1 (middle) and histamine (bottom). (**E**) AGR inst. ff. in response to 40 Hz IV neuron stimulation in saline (i), after blocking of H_2_ receptors with Cimetidine (ii) and after Cimetidine wash-out (iii). Recordings are from the same preparation and scaled identically. (**F**) Response of AGR inst. ff. to IV stimulation in saline (control stim) and after Cimetidine application for 3 preparations. Baseline refers to the AGR inst. ff. measured before IV stimulation. The decrease in AGR inst. ff. was significantly different from control stimulations. Paired t=test, p<0.01, N=3.

We have recently shown that the whole length of the central AGR axon is sensitive to modulation by the monoamine octopamine (OA, the invertebrate analog of norepinephrine), namely that AGR spike frequency increases independently of where OA is applied along the axon (Städele and Stein, 2015). To test whether histamine’s effects were restricted to the posterior *stn* or affected the whole axon trunk we also applied histamine specifically to anterior and middle parts of the *stn,* as well as to the *dgn*. In none of our experiments (N=3) did histamine cause a change in AGR firing frequency when applied to these *stn* parts.

To test whether histamine released from the IV neurons caused the diminishment of AGR’s firing frequency, we blocked the actions of histamine during IV neuron stimulation. We used cimetidine, an H_2_ receptor antagonist shown to effectively block IV neuron-mediated histaminergic effects in the STG (Christie et al., 2004). In specific, we stimulated the IV neurons in regular saline, observed the decrease in AGR firing frequency, applied cimetidine (5 mM) to AGR’s ectopic SIZ in the posterior *stn* and stimulated the IV neurons again. Figure 3E shows the response of AGR to IV neuron stimulation before and during cimetidine application. AGR’s response was significantly reduced in cimetidine (trace ii), indicating that histamine contributed to the IV neurons’ effect on AGR. The effect was reversible and the decrease in AGR firing frequency could be partly restored after cimetidine wash-out (trace iii). Across preparations, we found that in the presence of cimetidine the decrease in AGR firing frequency during IV stimulation was significantly reduced when compared to control stimulations (Fig. 3F, average AGR inst. ff. in control conditions: 2.23±0.76 Hz, cimetidine: 2.96±0.88 Hz, paired t-test, p<0.01, N=3). In summary, thus, our results demonstrate that the IV neuron co-transmitter histamine diminished AGR firing frequency, likely via H_2_ receptor activation. Since the decrease in AGR firing frequency was not fully suppressed when H_2_ receptors were blocked, other receptors may contribute as well.

### Descending projection neuron activity can relocate the site of spike initiation

We have recently shown that AGR can initiate action potentials everywhere along its axon trunk when excitability changes (Städele and Stein, 2015). Our results indicate that the IV neurons’ effect on AGR was targeted specifically at the ectopic spike initiation zone in the posterior *stn*. Thus, we predicted that spike initiation will remain at this particular spike initiation site until AGR firing frequency decreases below the intrinsic frequency of other axonal initiation sites. We monitored AGR’s spike activity extracellularly on the *dgn*, the anterior *stn* and the *son* and compared the delay between recording sites before, during and after IV stimulation. With the central spike initiation zone active, action potentials first occurred on the *stn* and then traveled antidromically towards the *dgn* and orthodromically towards the *son*. A relocation of the spike initiation site would cause a change in the relative spike timing between these recordings. Figure 4A shows the AGR spike occurrence of 70 action potentials for each recording site before, during and after IV neuron stimulation. We found that IV neuron activation did not change the initiation site during modest AGR frequency changes and that spike timing did not change during or after IV neuron stimulation (N=14).

**Figure 4.**
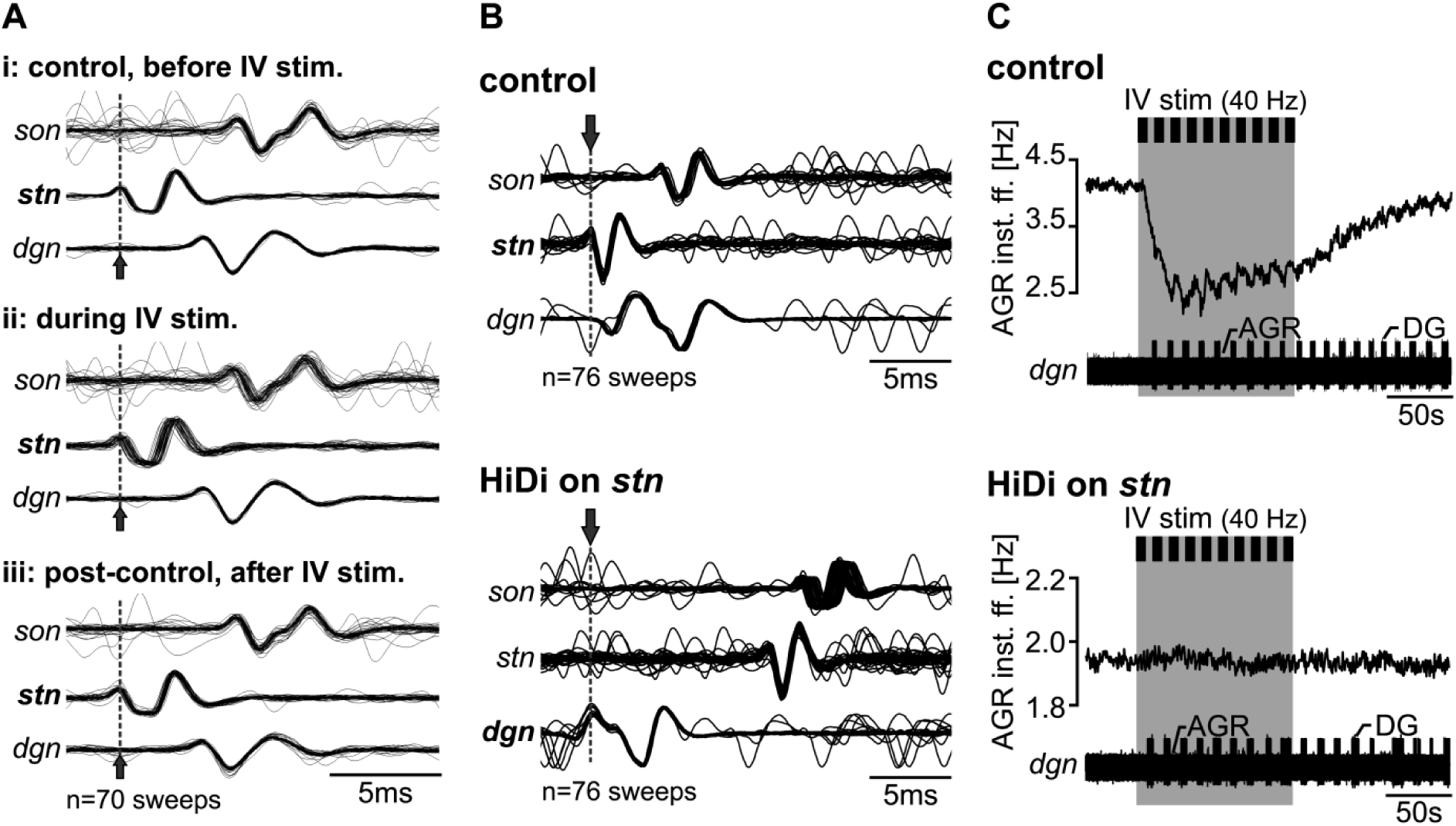
IV neuron stimulation does not relocate the site of spike initiation and specifically affects the *stn* spike initiation site. **(A)** Overlay of 70 AGR action potentials showing the spike occurrence at 3 extracellular recording sites before, during and after 40Hz IV neuron stimulation. Extracellular AGR spikes on the *stn* and *son* were aligned to the *dgn*. The site where the AGR action potential occurred first is indicated in bold. In control conditions (i) spikes first occurred on the *stn* and then traveled antidromically towards the *dgn* and orthodromically towards the *son*. IV neurons stimulation did not relocate the site of spike initiation since the delays between recording sites did not change during (ii) or after IV neuron stimulation (iii). **(B)** Change in conduction delays between *dgn*, *stn* and *son* before and after HiDi application to the posterior *stn*. HiDi application moved the site of spike initiation towards the periphery. In control condition (saline, top), action potentials first occurred on the *stn* recording. After HiDi application, action potentials first appear on the *dgn* and propagated orthodromically towards the *stn* and *son*. (**C**) AGR instantaneous firing frequency before and during IV neuron stimulation with spike activity generated at the eSIZ in the *stn* (top) and in the periphery (bottom, HiDi saline application to the posterior *stn*). The large unit on the *dgn* is the DG neuron. In HiDi, IV neuron stimulation did not affect AGR inst. ff. despite the fact that it elicited a strong and long-lasting gastric mill rhythm. Recordings from B and C are from the same preparation.

In several experiments (N=9 out of 45), however, the decrease in AGR firing frequency during IV neuron stimulation was so dramatic that the site of ectopic spike initiation switched to a different spike initiation site. In these cases spikes first occurred on the *dgn*. Such switches have previously been reported when the excitability at the *stn* spike initiation site is reduced. Daur et. al. (2009) showed that the *stn* SIZ competes with a spike initiation site in the *dgn* so that always the SIZ with the higher intrinsic firing frequency is active. Both *in vitro* and *in vivo*, the intrinsic firing frequency of the *dgn* SIZ is always lower than that of the *stn* SIZ. Our experiments show for the first time that descending neuromodulation can change this relation and allow for a switch of spike initiation zone: IV neuron activity reduced the intrinsic spike frequency of the *stn* SIZ to below that of the peripheral one, causing the switch of spike initiation site. Figure 5A and B shows the switch in AGR spike initiation during IV stimulation exemplarily for one preparation. Note the different time scales in Fig. 5A and B. In this particular example, IV neuron activation diminished AGR firing frequency by about 25% (from 2.8 to 2.1 Hz). Relatively quickly after the onset of the diminishment, AGR spike shape and amplitude on the *dgn* changed noticeably (arrows in Fig. 5B). As described previously (Daur et al., 2009) this change in AGR spike shape is an indication that the site of spike initiation moved to another location. When we compared the delay in spike occurrence on the different recordings, we found that spikes with negative amplitude were generated at the *stn* SIZ (Fig. 5C, trace i, *). In contrast, spikes with positive amplitude (**) were generated in the *dgn* (trace ii). In the example shown spike initiation switched back and forth between both SIZs, indicating that the intrinsic firing frequencies of both SIZs were approximately the same. The continuous switching affected spike timing dramatically and we found a strong jitter in spike timing (Fig. 5C, iii) when all action potentials with positive amplitude were taken into account.

**Figure 5.**
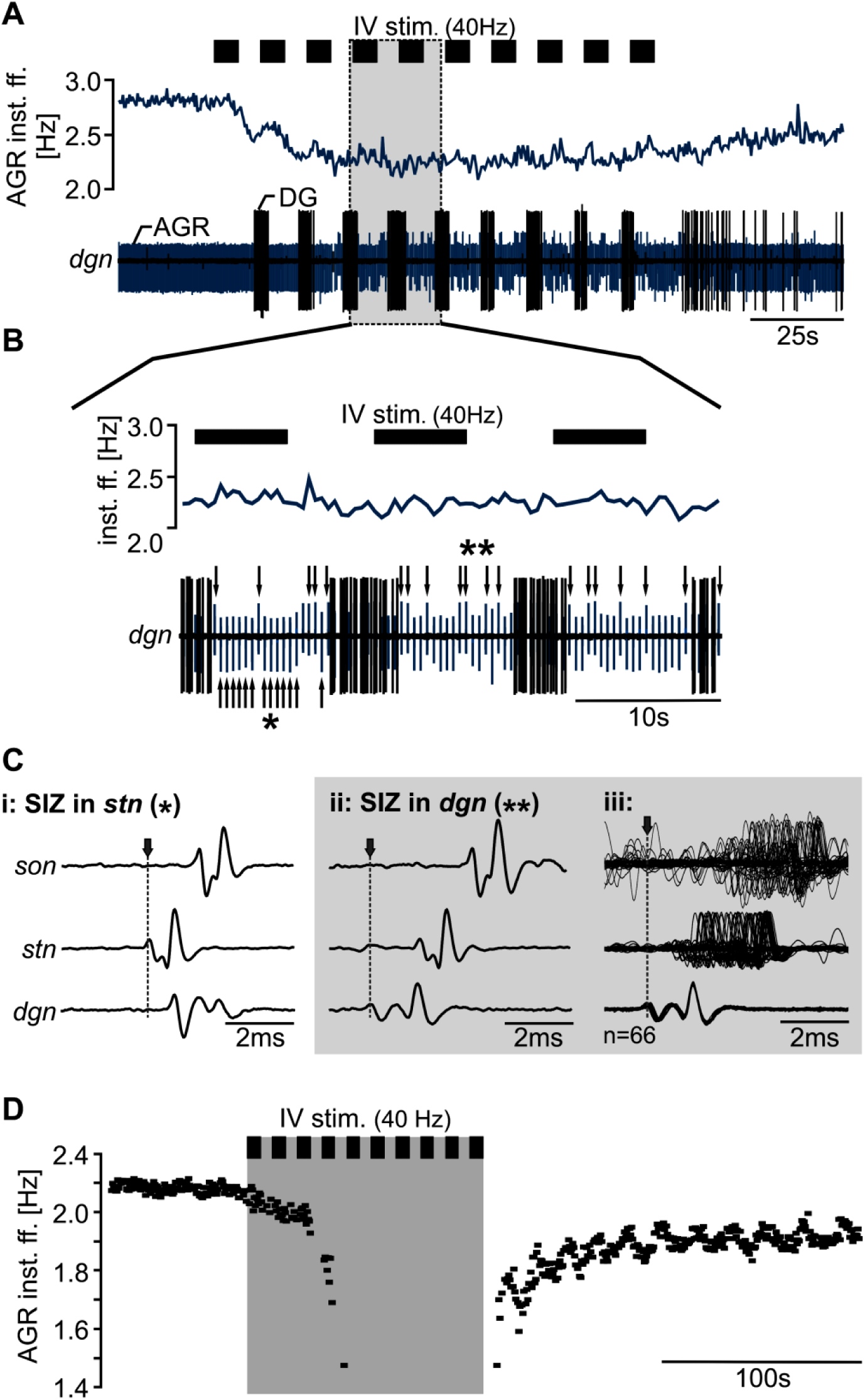
IV neuron stimulation can lead to a switch in the site of spike initiation. (**A**) Extracellular recording of the *dgn* showing that AGR spike amplitude changes during strong decrease of AGR instantaneous firing frequency (grey bar). (**B**) Magnification of the area marked in A. While AGR spikes on the *dgn* had similar shapes and amplitudes before IV neuron stimulation, spike amplitude continuously changed during IV stimulation. Arrows mark the changes in AGR amplitude. * highlights action potentials with large trough, ** those with large positive peaks. (**C**) Comparison of spike occurrence of AGR action potentials with large trough (i) and large positive peaks (ii) on the *dgn*, *stn* and *son*. Action potentials were aligned to the *dgn*. (i. ii) single sweeps. (iii) shows 66 action potentials with large positive peak. Action potentials with large trough (i) were generated at AGR’s central SIZ and appeared first on the *stn* whereas action potentials with large positive peak (ii) were initiated in the periphery and first appeared on the *dgn*. (iii) Jitter in spike timing of peripherally generated spikes. (**D**) Example for a complete stop in AGR spike initiation during IV stimulation. Shown is the change in AGR inst. ff. before, during (grey bar) and after repetitive 40 Hz IV stimulation for one preparation.

In conclusion, these results show that the IV neuron modulation can relocate the site of spike initiation by diminishing the AGR spike frequency at the *stn* SIZ. In some cases (N=4), IV stimulation even completely suppressed ectopic spike initiation (Fig. 5D).

To test whether histamine could be responsible for the switch in spike initiation site, we determined the site of spike initiation during focal application of histamine. Similar to IV neuron stimulation we found that in several experiments (N=5 out of 13) the decrease in AGR firing frequency was strong enough to switch the site of ectopic spike initiation to the *dgn* (arrows in Fig. 3C).

### Axonal modulation is spatially restricted

Since IV neuron stimulation shifted spike initiation to the *dgn*, we next tested whether this spike initiation site was similarly affected by descending modulation. We artificially moved the site of spike initiation away from the physiological location in the *stn* to the *dgn* via focal application of high-divalent saline (HiDi) to the posterior *stn*. HiDi is known to increase the spike threshold (Daur et al., 2009) and lead to a relocation of spike initiation to the *dgn* (Fig. 4B). This was also the case in our experiments. When we activated the IV neurons while spikes were initiated in the *dgn*, we found no further decrease in AGR firing frequency. Figure 4C shows the decrease in AGR firing frequency during 40 Hz IV neuron stimulation when spikes were elicited in the *stn* (normal saline) and when spikes were elicited in the *dgn* (HiDi application) for the same preparation. Although IV neuron stimulation elicited in both cases a strong and long-lasting gastric mill rhythm, AGR instantaneous firing frequency did not diminish when spikes were initiated in the *dgn*. Similarly as described by Städele and Stein (2015), the small gastric mill-timed oscillations of the AGR firing frequency were absent. We found this to be true for all preparations tested (N=5). In conclusion, thus, descending modulation specifically targeted the *stn* spike initiation site, allowing a direct regulation of ectopic spike frequency at this site. In contrast, the *dgn* spike initiation site was unaffected by descending modulation.

### Axon propagation dynamics are not influenced by descending modulation

Axons are often seen as faithful conduits of action potentials. However, there are numerous reports demonstrating that action potential transmission and thus information content can be modified on its way to postsynaptic targets (REF). Ectopic spike initiation elicited by neuromodulators like dopamine, serotonin or octopamine, for example, can add additional action potentials to the ones generated at the axon hillock (Meyrand et al., 1992; Bucher et al., 2003; Goaillard et al., 2004; Städele and Stein, 2015). Neuromodulators can also affect the dynamics of action potential propagation and affect spike frequency in a history-dependent way (Panzeri et al., 2001; Petersen et al., 2001; Lang et al., 2006; Ballo and Bucher, 2009). We have shown above that the IV neurons regulate ectopic action potential frequency. We thus tested whether this regulation also results in a modification of en-passant action potentials as they pass the site of modulation. Such modulation could result in a change in conduction dynamics and thus a change in the temporal structure of spike discharges or even spike failures.

We activated AGR in the periphery and measured the delays with which action potentials occurred on the *stn* and *son* before and during IV neuron modulation. In specific, we stimulated AGR extracellularly on the anterior gastric nerve (*agn*), a nerve that exclusively contains the AGR axon and innervates the muscles in the periphery. The *agn* was stimulated with 5 consecutive trains, each with 28 pulse and 15 Hz stimulation frequency, which approximately corresponds to AGR’s *in vivo* activity (Smarandache and Stein, 2007; Daur et al., 2009; Daur et al., 2012). Figure 6A shows the occurrence of action potentials on the *stn* and *son* before (control) and during IV modulation in one experiment. Action potentials were aligned to the stimulus and plotted on top of each other so that the first spike appears at the bottom and the last one at the top. For better comparison, spike times were extracted and plotted as a function of delay to the *agn* stimulation (Fig. 6B). We found that the AGR conduction velocity was history-dependent even in control in that it initially decreased and then increased.. When we compared the temporal occurrence of each AGR spike during IV neuron stimulation to that in control we found no change. Figure 6C shows the difference in spike delay (=Δ delay) on the *stn* and *son* before and during IV neuron modulation for 6 experiments. On average, we found that Δ delay for each spike was not significantly different from 0 (one sided t-test, p>0.5, N=6). Thus, IV neuron modulation of AGR spike initiation did not influence the temporal coding of en-passant action potentials.

**Figure 6.**
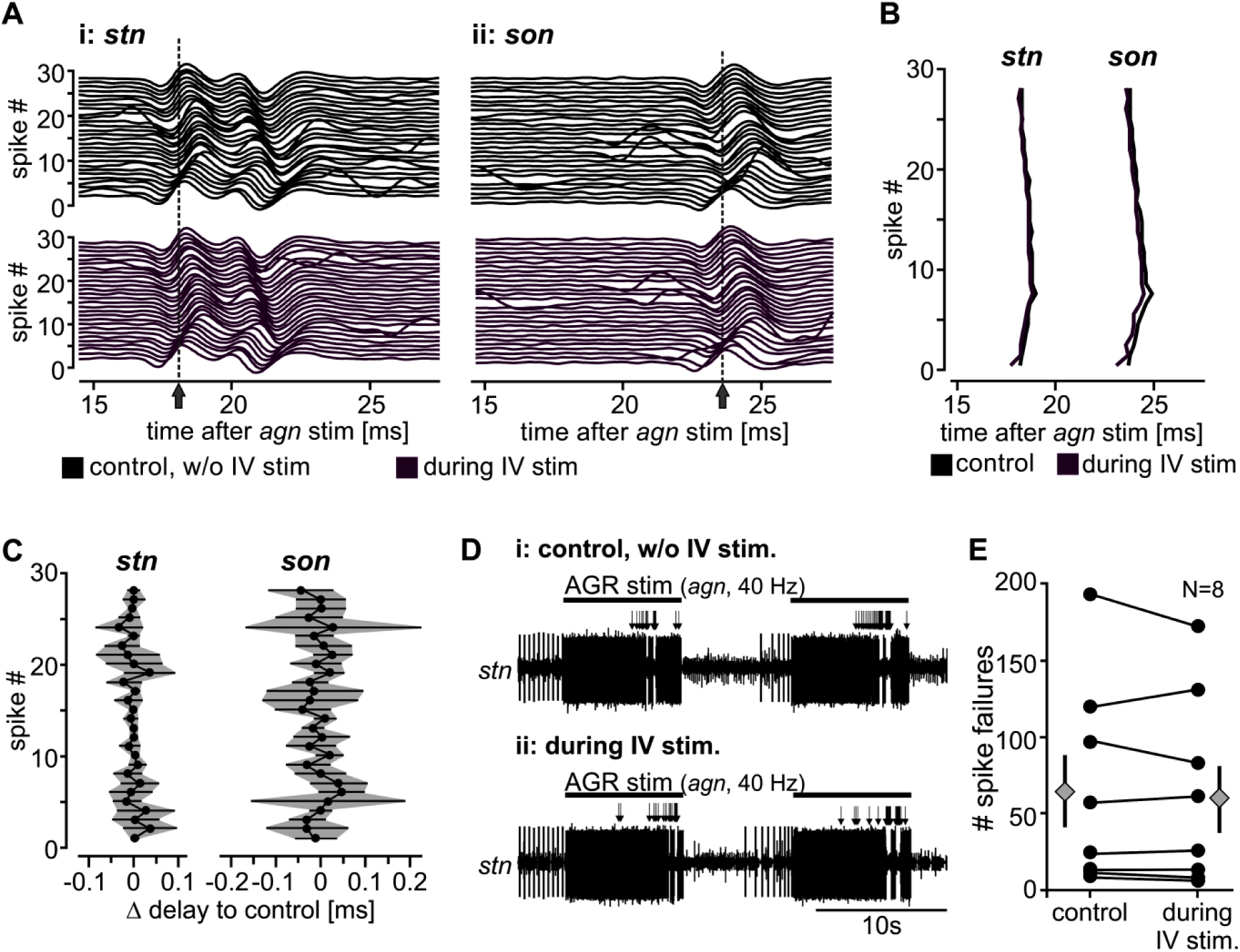
IV neuron modulation does not affect en-passant information. (**A**) Single burst of peripherally initiated AGR action potentials in response to extracellular stimulation of the *agn* with 28 consecutive pulses and 15 Hz stimulation frequency. Shown are spike occurrences on *stn* (i) and *son* (ii) before (black) and during IV neuron stimulation (purple). Action potentials were aligned to the stimulus and plotted on top of each other so that the first spike occurs at the bottom. The arrow marks the time of the first peak of the first action potential. (**B**) Comparison of spike occurrence for the example shown in A. Spike times were extracted and plotted as a function of delay to the *agn* stimulation. (**C**) Analysis of the difference in spike delay (=Δ delay) on the *stn* and *son* before (control) and during IV neuron modulation for 6 experiments. The delay was calculated by subtracting the time when a given action potential occurred on the extracellular recording site during IV neuron stimulation from control values. Shown are means ± SD (grey area). (**D**) Example extracellular recording of the *stn* showing spike failures of peripherally activated AGR action potentials before and during IV neuron modulation. Recordings are from the same preparation. AGR was activated via extracellular repetitive *agn* stimulation (40 Hz, 9 sec train/intertrain duration). Arrows highlight areas where spike failures occurred. (**E**) Analysis of the number of spike failures during repetitive *agn* stimulation with 40 Hz before (control) and during IV neuron modulation. Circles represent data from single experiments; diamonds represent the mean ± SEM, N=6.

Moderate or high frequency stimulation (10-50 Hz) of axons can lead to propagation failures (Krnjevic and Miledi, 1959; Grossman et al., 1979) so that action potentials fail to propagate along the axon and information is ‘deleted’ before it reaches the postsynaptic target. Neuromodulators that act on axon can cause excitability changes and may thus alter the rate of conduction failures (Debanne, 2004). We tested whether the rate of propagation failures and the maximum conduction frequency in AGR is affected by the IV neurons. We stimulated the *agn* with 10 trains of 10 to 50 Hz (10 Hz steps) for 9 seconds and compared the number of spikes passing through the site of modulation before and during IV neuron activation. 50 Hz simulation reliably caused spike failures of >50% even in control conditions (without IV neuron modulation), indicating that AGR’s maximum firing frequency is limited to lower frequencies. Stimulation frequencies between 10 and 30 Hz did not result in any spike failures (neither in control nor during IV neuron modulation). At 40 Hz (arrows in Fig. 6 D) spike failures started to occur both in control and during IV neuron stimulation. On average, however, we found no significant difference between the number of spike failures during IV neuron modulation when compared to control conditions (Fig. 6E, paired t-test, p=0.5, N=6). Thus, the maximum transmission frequency of the AGR axon appeared to be unaffected by IV neuron modulation. Taken together, our results demonstrate that descending modulation of axonal spike initiation did not affect axonal propagation dynamics.

## Discussion

### Descending control of sensory spike initiation

Many vital behaviors such as breathing, swallowing, and chewing, as well as locomotion and saccadic eye movements are driven by central pattern generating circuits that produce rhythmic output patterns (Marder and Calabrese, 1996; Grillner et al., 2005; Dickinson, 2006; Gordon and Whelan, 2006; Isa and Sparks, 2006; Kiehn, 2006; Katz and Hooper, 2007; Chevallier et al., 2008; Doi and Ramirez, 2008; Berkowitz et al., 2010; El Manira et al., 2010; Büschges et al., 2011; Harris-Warrick, 2011; Marder, 2012). Some behaviors driven by CPGs are continuous and stereotypic (Calabrese and De Schutter, 1992; Cleland and Selverston, 1998; Norris et al., 2011; Roffman et al., 2012) while others are episodic and flexible (Katz, 2009; Selverston et al., 2009; Büschges et al., 2011). This flexibility is often the result of numerous influences affecting CPG circuits, such as hormonal or neurally released neuromodulators, sensory pathways and descending projections.

Descending modulatory projection neurons, for example, modify CPG output on almost every level, including short and long-term modifications of synapses and ionic conductances in circuit neurons as well as other projection neurons (McClellan, 1984; Stein, 2009; Shaw et al., 2010; Blitz and Nusbaum, 2011; von Philipsborn et al., 2011; Thiele et al., 2014; Borgmann and Buschges, 2015). Modulation results in an immense plasticity of motor network output through dynamic adaptation of existing pattern generating circuits. Descending neuromodulatory control is thus fundamentally involved in the process how nervous systems select distinct behaviors.

To our knowledge, this study is the first that demonstrates two distinct actions of descending projection neurons, namely that spike initiation in axons is under extrinsic neuromodulatory control and that descending projection neurons directly modulate sensory neurons. So far there were no indications that ectopic spike initiation in sensory neurons can be directly regulated by descending neurons. While neuromodulators such as neuropeptides and monoamines have been shown to affect both muscle contraction and sensory activity in the periphery (Birmingham et al., 1999; Birmingham, 2001), there is no evidence of the source of these modulators. In many cases, assumptions about modulator sources stem from the concentration threshold for bath-applied neuromodulators. If the concentration is low, it is generally assumed that they reach the periphery as circulating hormones (Meyrand et al., 1992; Bucher et al., 2003; Goaillard et al., 2004). If modulators were released from descending neurons, however, this could allow CPGs to interfere with sensory spike activity, at least in principle. CPGs provide ascending feedback onto descending modulatory neurons that control CPG activity, but also release neuromodulatory substances in a paracrine fashion (Antri et al., 2009; Buchanan, 2011; Blitz and Nusbaum, 2012). If these neuromodulators reach the periphery, muscle contraction and sensory activity may be affected, allowing CPGs to indirectly control sensory activity.

Sensory pathways have been shown to interact on many levels with CPGs and their descending inputs to dynamically adjust CPG output in response to changing internal and external conditions (Bässler, 1986; Wolf, 1991; Prochazka et al., 2002; Beenhakker et al., 2004; Büschges, 2005; Doi and Ramirez, 2008; DeLong and Nusbaum, 2010; Büschges et al., 2011; Stein, 2014). While most sensory influences are feed-forward in that the sensory neurons inform the CNS of changes in the environment or the body, there is also feedback from the CNS to the afferent pathways. In particular, gating of sensory and sensory-related information in central pathways via presynaptic inhibition is common in many systems (el Manira and Clarac, 1994; Coleman et al., 1995; Sauer et al., 1997; Stein and Schmitz, 1999; Beenhakker et al., 2005; Beenhakker et al., 2007; Barriere et al., 2008; DeLong et al., 2009; Bardoni et al., 2013). In locusts, for example, the sensory feedback from the femoral chordotonal organ, which measures the position of the femur-tibia joint during walking, is gated with the phase of the walking motor pattern, disabling feedback during certain phases of this pattern (Wolf and Burrows, 1995). The underlying mechanism is a phase-dependent presynaptic inhibition of the sensory terminals and a concomitant decrease in transmitter release without affecting sensory activity and action potential propagation to the terminals. While the central pathways that elicit presynaptic inhibition are still somewhat nebulous, it has been demonstrated that proprioceptors can presynaptically inhibit the terminals of neurons from other proprioceptive sense organs (Stein and Schmitz, 1999, Clarac and Cattaert, 1996; Cattaert et al., 1999; Barriere et al., 2008). Since proprioceptors typically show phasic activity, this could contribute to the gating of sensory feedback at specific phases of the motor pattern.

Gating of sensory information often results in a diminishment or even complete block of afferent spike propagation at the terminal (Burrows and Matheson, 1994; Clarac and Cattaert, 1996; Schmitz and Stein, 2000; HYPERLINK \l “bookmark79” Margrie et al., 2001; Xiong and Chen, 2002; Evans et al., 2003; Barriere et al., 2008; Lee et al., 2012). In contrast, our results show that afferent spike transmission in AGR was not gated by IV neurons. Modulation of ectopic spike initiation had no influence on axon dynamics and action potential propagation dynamics of en-passant action potentials (Fig. 6). In specific, there was no influence of IV neuron activity on action potential propagation delay and history-dependence. This occurred despite the fact that modulator actions on axons have been shown to determine propagation dynamics (Panzeri et al., 2001; Ballo and Bucher, 2009; Ballo et al., 2010; Ballo et al., 2012). However, modulatory influences on propagation dynamics are more effective if they are exerted along a large stretch of the axon (Ballo and Bucher, 2009). This was not the case in our experiments as the spike initiation zone is a rather restricted locus on the axon. There was also no effect on the maximum frequency, i.e. action potential conduction seemed unaffected.

In summary, thus, the modulation of axonal spike initiation by descending projection neurons represents a convenient way of altering sensory spike activity without affecting afferent information transfer. Since most motor networks are under descending modulatory control and receive sensory feedback, direct neuronal control of sensory spike initiation may be a common and understudied principle for activity-dependent regulation of sensory activity.

### Mechanism of modulation

How can descending modulatory neurons affect spike initiation without altering spike propagation dynamics? The IV neurons contain at least 2 co-transmitters (histamine and FMRF-like peptide F1, Christie et al., 2004; Stein et al., 2007). Our data show that while both can affect AGR, only histamine caused a decrease in AGR firing rate (Fig. 3C). We were able to partially block the histamine effect with the competitive H_2_ receptor antagonist cimetidine (Fig. 3E). Histamine, just like other biogenic amines, has been shown to mediate a plethora of different actions on neurons. It is known to be involved in thermoregulation, feeding rhythms, sensory and motor control, and circadian control of sleep, metabolic rates and appetite (summarized in Haas et al., 2008). Histamine can either act via metabotropic G-protein coupled receptors (H_1_-H_4_) that mediate an increase in intracellular cAMP or via ionotropic receptors that activate chloride conductances. The latter can be found in the hypothalamus (Hatton and Yang, 2001) and thalamus (Lee et al., 2004), olfactory neurons of the spiny lobster (McClintock and Ache, 1989), neurosecretory cells in crayfish (Cebada and Garcia, 2007), motor neurons of the lobster cardiac ganglion (Hashemzadeh-Gargari and Freschi, 1992) and the antennal lobe in honey bee (Sachse et al., 2006). Albeit the overall effect on neural networks is mostly inhibitory, this inhibition is typically indirect and mediated via excitation of inhibitory interneurons (Haas et al., 2008). In the STNS, in contrast, all hitherto described histaminergic actions are inhibitory (Claiborne and Selverston, 1984; Pulver et al., 2003; Christie et al., 2004). Histamine elicits a strong and fast hyperpolarization of the pyloric pacemaker neurons in the STG that subsides a few seconds after the end of the IV neuron stimulation (Christie et al., 2004). Since the histaminergic effects on AGR were much slower and maintained for much longer, the kinetics seem distinct from histaminergic actions exerted on STG motor neurons.

What mechanisms could be responsible for the observed decrease in ectopic spike initiation during IV neuron activity? Potential mechanisms include a reduction in depolarizing ionic conductances or activation of hyperpolarizing conductances (e.g. chloride or potassium) by histamine. We found that blocking H_2_ receptors with cimetidine substantially diminished the IV neuron mediated effect (Fig. 3E). H_2_ receptors have been shown to interact with ionic conductance such as I_K(Ca)_ (Haas and Konnerth, 1983; Pedarzani and Storm, 1993) and I_h_ (McCormick and Williamson, 1991; Pedarzani and Storm, 1995). While I_h_ is present in AGR (Daur et al., 2012), I_K(Ca)_ has not been identified yet. In any case, a histamine-induced reduction of either conductance or a shift in their voltage- or calcium-dependence could alter cell resting potential and spike activity. For example, Ballo et al. (2010) show that axonal I_h_ depolarizes the resting membrane potential in the axon and causes ectopic spiking. However, due to its depolarizing effect, I_h_ also affects action potential propagation. In specific, I_h_ influences the slow hyperpolarization (sAHP) of action potentials (Grafe et al., 1997; Soleng et al., 2003; Baginskas et al., 2009). The relative contribution of I_h_ determines the degree to which an axon hyperpolarizes during repetitive activity (Kiernan et al., 2004). The sAHP, in turn, has been implicated in a general slowing of action potential propagation (Bostock and Grafe, 1985; Grafe et al., 1997; Moalem-Taylor et al., 2007; Ballo et al., 2010; Ballo et al., 2012). The fact that in our experiments action potential propagation during IV neuron modulation did not change (Fig. 6) indicates that if H_2_ receptors modulate I_h_ or another conductance in AGR, this effect is spatially restricted to the ectopic spike initiation zone.

We noted that histamine influenced AGR only at rather high concentrations (10 mM, Fig. 3C). The high concentration needed in our experiments might be explained in part by the fact that all neuromodulators were applied to the non-desheathed nervous system (see Material and Methods). The reason for this was that removal of the sheath impaired the IV neuron-mediated influence on AGR, similarly to previously described effects by Goldsmith et al (2014). The high concentrations needed could indicate that histamine is unable to fully penetrate the nerve sheath. This further supports the idea that histamine reaches the AGR axon only via chemical synaptic transmission rather than as circulating hormone. Hormone concentrations are typically very low in contrast to neurally released transmitters. If the sheath indeed interferes with histamine actions then low hormonal concentrations will not be sufficient to affect AGR.

### Functional implications

The result of the AGR modulation is a change in its tonic spike frequency. Changes of a few Hertz in tonic activity modify ongoing gastric mill rhythms (Daur et al., 2009). Magnitude and timing of the gastric mill motor pattern correlate with the tonic spike activity of the muscle receptor, making the state of the system and ongoing activity important contributors to stimulus-induced changes in motor output. Interestingly, the effect on AGR was specific to the IV neurons and not elicited by a mechanosensory pathway (VCN) that also elicits gastric mill rhythms (Fig. 1). In fact, VCN elicits its version of the rhythm via the same set of descending projection neurons (MCN1 and CPN2) as the IV neurons (Beenhakker and Nusbaum, 2004; Hedrich et al., 2009), indicating that not the mere presence of descending modulation but its distinct type determines AGR axon modulation. The differential response to different pathways could represent an understudied form of neural plasticity in that not only the ongoing motor activity but even sensory feedback created by it is differentially affected by distinct descending pathways.

## Acknowledgements

We would like to thank Dr. Lingjun Li for providing us with the Orcokinins used in this study. This work was supported by grants from the German Research Foundation (DFG STE 937/9- 1), National Science Foundation (NSF IOS 1354932), Illinois State University and The German Academic Exchange Service (DAAD).

## References

Antri M, Fenelon K, Dubuc R (2009) The contribution of synaptic inputs to sustained depolarizations in reticulospinal neurons. J Neurosci 29:1140–1151.

Ausborn J, Stein W, Wolf H (2007) Frequency Control of Motor Patterning by Negative Sensory Feedback. J Neurosci 27:9319–9328.

Baginskas A, Palani D, Chiu K, Raastad M (2009) The H-current secures action potential transmission at high frequencies in rat cerebellar parallel fibers. Eur J Neurosci 29:87–96.

Ballo AW, Bucher D (2009) Complex intrinsic membrane properties and dopamine shape spiking activity in a motor axon. J Neurosci 29:5062–5074.

Ballo AW, Nadim F, Bucher D (2012) Dopamine modulation of Ih improves temporal fidelity of spike propagation in an unmyelinated axon. J Neurosci 32:5106–5119.

Ballo AW, Keene JC, Troy PJ, Goeritz ML, Nadim F, Bucher D (2010) Dopamine Modulates Ih in a Motor Axon. J Neurosci 30:8425–8434.

Bardoni R, Takazawa T, Tong CK, Choudhury P, Scherrer G, Macdermott AB (2013) Pre- and postsynaptic inhibitory control in the spinal cord dorsal horn. Annals of the New York Academy of Sciences 1279:90–96.

Barriere G, Simmers J, Combes D (2008) Multiple mechanisms for integrating proprioceptive inputs that converge on the same motor pattern-generating network. J Neurosci 28:8810–8820.

Bässler U (1986) On the definition of central pattern generator and its sensory control. Biol Cybern 54:65–69.

Beenhakker MP, Nusbaum MP (2004) Mechanosensory activation of a motor circuit by coactivation of two projection neurons. J Neurosci 24:6741–6750.

Beenhakker MP, Blitz DM, Nusbaum MP (2004) Long-lasting activation of rhythmic neuronal activity by a novel mechanosensory system in the crustacean stomatogastric nervous system. J Neurophysiol 91:78–91.

Beenhakker MP, Kirby MS, Nusbaum MP (2007) Mechanosensory gating of proprioceptor input to modulatory projection neurons. J Neurosci 27:14308–14316.

Beenhakker MP, DeLong ND, Saideman SR, Nadim F, Nusbaum MP (2005) Proprioceptor regulation of motor circuit activity by presynaptic inhibition of a modulatory projection neuron. J Neurosci 25:8794–8806.

Berkowitz A, Roberts A, Soffe SR (2010) Roles for multifunctional and specialized spinal interneurons during motor pattern generation in tadpoles, zebrafish larvae, and turtles. Front Behav Neurosci 4:36.

Birmingham JT (2001) Increasing sensor flexibility through neuromodulation. Biol Bull 200:206–210.

Birmingham JT, Szuts ZB, Abbott LF, Marder E (1999) Encoding of muscle movement on two time scales by a sensory neuron that switches between spiking and bursting modes. J Neurophysiol 82:2786–2797.

Birmingham JT, Billimoria CP, DeKlotz TR, Stewart RA, Marder E (2003) Differential and history-dependent modulation of a stretch receptor in the stomatogastric system of the crab, *Cancer borealis*. J Neurophysiol 90:3608–3616.

Blitz DM, Nusbaum MP (2011) Neural circuit flexibility in a small sensorimotor system. Curr Opin Neurobiol 21:544–552.

Blitz DM, Nusbaum MP (2012) Modulation of circuit feedback specifies motor circuit output. J Neurosci 32:9182–9193.

Borgmann A, Buschges A (2015) Insect motor control: methodological advances, descending control and inter-leg coordination on the move. Curr Opin Neurobiol 33:8–15.

Bostock H, Grafe P (1985) Activity-dependent excitability changes in normal and demyelinated rat spinal root axons. J Physiol 365:239–257.

Buchanan JT (2011) Spinal locomotor inputs to individually identified reticulospinal neurons in the lamprey. J Neurophysiol 106:2346–2357.

Bucher D, Goaillard J-M (2011) Beyond faithful conduction: short-term dynamics, neuromodulation, and long-term regulation of spike propagation in the axon. Prog Neurobiol 94:307–346.

Bucher D, Thirumalai V, Marder E (2003) Axonal dopamine receptors activate peripheral spike initiation in a stomatogastric motor neuron. J Neurosci 23:6866–6875.

Burrows M, Matheson T (1994) A presynaptic gain control mechanism among sensory neurons of a locust leg proprioceptor. J Neurosci 14:272–282.

Büschges A (2005) Sensory control and organization of neural networks mediating coordination of multisegmental organs for locomotion. J Neurophysiol 93:1127–1135.

Büschges A, Scholz H, El Manira A (2011) New moves in motor control. Curr Biol 21:R513–524.

Calabrese RL, De Schutter E (1992) Motor-pattern-generating networks in invertebrates: modeling our way toward understanding. Trends Neurosci 15:439–445.

Cattaert D, El Manira A, Bevengut M (1999) Presynaptic inhibition and antidromic discharges in crayfish primary afferents. J Physiol Paris 93:349–358.

Cebada J, Garcia U (2007) Histamine operates Cl-gated channels in crayfish neurosecretory cells. J Exp Biol 210:3962–3969.

Chevallier S, Jan Ijspeert A, Ryczko D, Nagy F, Cabelguen JM (2008) Organisation of the spinal central pattern generators for locomotion in the salamander: biology and modelling. Brain Res Rev 57:147–161.

Christie AE, Stein W, Quinlan JE, Beenhakker MP, Marder E, Nusbaum MP (2004) Actions of a histaminergic/peptidergic projection neuron on rhythmic motor patterns in the stomatogastric nervous system of the crab *Cancer borealis*. J Comp Neurol 469:153–169.

Claiborne BJ, Selverston AI (1984) Histamine as a neurotransmitter in the stomatogastric nervous system of the spiny lobster. J Neurosci 4:708–721.

Clarac F, Cattaert D (1996) Invertebrate presynaptic inhibition and motor control. Exp Brain Res 112:163–180.

Cleland TA, Selverston AI (1998) Inhibitory glutamate receptor channels in cultured lobster stomatogastric neurons. J Neurophysiol 79:3189–3196.

Coleman MJ, Meyrand P, Nusbaum MP (1995) A switch between two modes of synaptic transmission mediated by presynaptic inhibition. Nature 378:502–505.

Combes D, Simmers J, Moulins M (1995) Structural and functional characterization of a muscle tendon proprioceptor in lobster. J Comp Neurol 363:221–234.

Cropper EC, Evans CG, Jing J, Klein A, Proekt A, Romero A, Rosen SC (2004) Regulation of afferent transmission in the feeding circuitry of Aplysia. Acta Biol Hung 55:211–220.

Daur N, Nadim F, Stein W (2009) Regulation of motor patterns by the central spike-initiation zone of a sensory neuron. Eur J Neurosci 30:808–822.

Daur N, Diehl F, Mader W, Stein W (2012) The stomatogastric nervous system as a model for studying sensorimotor interactions in real-time closed-loop conditions. Front Comput Neurosci 6.

Debanne D (2004) Information processing in the axon. Nat Rev Neurosci 5:304–316.

Delcomyn F (1980) Neural basis of rhythmic behavior in animals. Science 210:492–498.

DeLong ND, Nusbaum MP (2010) Hormonal modulation of sensorimotor integration. J Neurosci 30:2418–2427.

DeLong ND, Beenhakker MP, Nusbaum MP (2009) Presynaptic inhibition selectively weakens peptidergic cotransmission in a small motor system. J Neurophysiol 102:3492–3504.

Dickinson PS (2006) Neuromodulation of central pattern generators in invertebrates and vertebrates. Curr Opin Neurobiol 16:604–614.

Doi A, Ramirez JM (2008) Neuromodulation and the orchestration of the respiratory rhythm. Respir Physiol Neurobiol 164:96–104.

El Manira A, Clarac F (1994) Presynaptic inhibition is mediated by histamine and GABA in the crustacean escape reaction. J Neurophysiol 71:1088–1095.

El Manira A, Kyriakatos A, Nanou E (2010) Beyond connectivity of locomotor circuitry-ionic and modulatory mechanisms. Prog Brain Res 187:99–110.

Evans CG, Jing J, Rosen SC, Cropper EC (2003) Regulation of Spike Initiation and Propagation in an Aplysia Sensory Neuron: Gating-In via Central Depolarization. J Neurosci 23:2920–2931.

Goaillard JM, Schulz DJ, Kilman VL, Marder E (2004) Octopamine modulates the axons of modulatory projection neurons. J Neurosci 24:7063–7073.

Goeritz ML, Bowers MR, Slepian B, Marder E (2013) Neuropilar projections of the anterior gastric receptor neuron in the stomatogastric ganglion of the Jonah crab, *Cancer borealis*. PLoS One 8:e79306.

Goldsmith CJ, Städele C, Stein W (2014) Optical imaging of neuronal activity and visualization of fine neural structures in non-desheathed nervous systems. PLoS One 9:e103459.

Gordon IT, Whelan PJ (2006) Deciphering the organization and modulation of spinal locomotor central pattern generators. J Exp Biol 209:2007–2014.

Grafe P, Quasthoff S, Grosskreutz J, Alzheimer C (1997) Function of the hyperpolarization-activated inward rectification in nonmyelinated peripheral rat and human axons. J Neurophysiol 77:421–426.

Grillner S (2009) Networks in Motion. Brain Res Rev:1–274.

Grillner S, Markram H, De Schutter E, Silberberg G, LeBeau FE (2005) Microcircuits in action-from CPGs to neocortex. Trends Neurosci 28:525–533.

Grossman Y, Parnas I, Spira ME (1979) Differential conduction block in branches of a bifurcating axon. J Physiol 295:283–305.

Haas HL, Konnerth A (1983) Histamine and noradrenaline decrease calcium-activated potassium conductance in hippocampal pyramidal cells. Nature 302:432–434.

Haas HL, Sergeeva OA, Selbach O (2008) Histamine in the nervous system. Physiol Rev 88:1183–1241.

Harris-Warrick RM (2011) Neuromodulation and flexibility in Central Pattern Generator networks. Curr Opin Neurobiol 21:685–692.

Hashemzadeh-Gargari H, Freschi JE (1992) Histamine activates chloride conductance in motor neurons of the lobster cardiac ganglion. J Neurophysiol 68:9–15.

Hatton GI, Yang QZ (2001) Ionotropic histamine receptors and H2 receptors modulate supraoptic oxytocin neuronal excitability and dye coupling. J Neurosci 21:2974–2982.

Hedrich UB, Stein W (2008) Characterization of a descending pathway: activation and effects on motor patterns in the brachyuran crustacean stomatogastric nervous system. J Exp Biol 211:2624–2637.

Hedrich UB, Smarandache CR, Stein W (2009) Differential activation of projection neurons by two sensory pathways contributes to motor pattern selection. J Neurophysiol 102:2866–2879.

Hedrich UB, Diehl F, Stein W (2011) Gastric and pyloric motor pattern control by a modulatory projection neuron in the intact crab *Cancer pagurus*. J Neurophysiol 105:1671–1680.

Isa T, Sparks DL (2006) Microcircuit of the Superior Colliculus. A Neuronal Machine that Determines Timing and Endpoint of Saccadic Eye Movements. In: Microcircuits: The Interface between Neurons and Global Brain Function (Grillner S, Graybiel AM, eds), pp 5–34.

Katz PS, Frost WN (1996) Intrinsic neuromodulation: altering neuronal circuits from within. Trends Neurosci 19:54–61.

Katz PS, Hooper SL (2007) Invertebrate central pattern generators. Cold Spring Harbor Press 49:251.

Kiehn O (2006) Locomotor circuits in the mammalian spinal cord. Annu Rev Neurosci 29:279–306.

Kiernan MC, Lin CS, Burke D (2004) Differences in activity-dependent hyperpolarization in human sensory and motor axons. J Physiol 558:341–349.

Krnjevic K, Miledi R (1959) Presynaptic failure of neuromuscular propagation in rats. J Physiol 149:1–22.

Lang PM, Moalem-Taylor G, Tracey DJ, Bostock H, Grafe P (2006) Activity-dependent modulation of axonal excitability in unmyelinated peripheral rat nerve fibers by the 5-HT(3) serotonin receptor. J Neurophysiol 96:2963–2971.

Lee AH, Megalou EV, Wang J, Frost WN (2012) Axonal conduction block as a novel mechanism of prepulse inhibition. J Neurosci 32:15262–15270.

Lee KH, Broberger C, Kim U, McCormick DA (2004) Histamine modulates thalamocortical activity by activating a chloride conductance in ferret perigeniculate neurons. Proc Natl Acad Sci USA 101:6716–6721.

Lena C, Changeux JP, Mulle C (1993) Evidence for “preterminal” nicotinic receptors on GABAergic axons in the rat interpeduncular nucleus. J Neurosci 13:2680–2688.

Li L, Pulver SR, Kelley WP, Thirumalai V, Sweedler JV, Marder E (2002) Orcokinin peptides in developing and adult crustacean stomatogastric nervous systems and pericardial organs. J Comp Neurol 444:227–244.

Marder E (2012) Neuromodulation of neuronal circuits: back to the future. Neuron 76:1–11.

Marder E, Calabrese RL (1996) Principles of rhythmic motor pattern generation. Physiol Rev 76:687–717.

Marder E, Bucher D (2001) Central pattern generators and the control of rhythmic movements. Curr Biol 11:R986–996.

Margrie TW, Sakmann B, Urban NN (2001) Action potential propagation in mitral cell lateral dendrites is decremental and controls recurrent and lateral inhibition in the mammalian olfactory bulb. Proc Natl Acad Sci USA 98:319–324.

McClellan AD (1984) Descending control and sensory gating of ‘fictive’ swimming and turning responses elicited in an in vitro preparation of the lamprey brainstem/spinal cord. Brain Res 302:151–162.

McClintock TS, Ache BW (1989) Histamine directly gates a chloride channel in lobster olfactory receptor neurons. Proc Natl Acad Sci USA 86:8137–8141.

McCormick DA, Williamson A (1991) Modulation of neuronal firing mode in cat and guinea pig LGNd by histamine: possible cellular mechanisms of histaminergic control of arousal. J Neurosci 11:3188–3199.

Meyrand P, Weimann JM, Marder E (1992) Multiple axonal spike initiation zones in a motor neuron: serotonin activation. J Neurosci 12:2803–2812.

Mitchell GS, Johnson SM (2003) Neuroplasticity in respiratory motor control. J Appl Physiol (1985) 94:358–374.

Moalem-Taylor G, Lang PM, Tracey DJ, Grafe P (2007) Post-spike excitability indicates changes in membrane potential of isolated C-fibers. Muscle Nerve 36:172–182.

Nadim F, Bucher D (2014) Neuromodulation of neurons and synapses. Curr Opin Neurobiol 29C:48–56.

Norris BJ, Wenning A, Wright TM, Calabrese RL (2011) Constancy and variability in the output of a central pattern generator. J Neurosci 31:4663–4674.

Nusbaum MP, Beenhakker MP (2002) A small-systems approach to motor pattern generation. Nature 417:343–350.

Panzeri S, Petersen RS, Schultz SR, Lebedev M, Diamond ME (2001) The role of spike timing in the coding of stimulus location in rat somatosensory cortex. Neuron 29:769–777.

Pedarzani P, Storm JF (1993) PKA mediates the effects of monoamine transmitters on the K+ current underlying the slow spike frequency adaptation in hippocampal neurons. Neuron 11:1023–1035.

Pedarzani P, Storm JF (1995) Protein kinase A-independent modulation of ion channels in the brain by cyclic AMP. Proc Natl Acad Sci USA 92:11716–11720.

Petersen RS, Panzeri S, Diamond ME (2001) Population coding of stimulus location in rat somatosensory cortex. Neuron 32:503–514.

Pinault D (1995) Backpropagation of action potentials generated at ectopic axonal loci: hypothesis that axon terminals integrate local environmental signals. Brain Res Brain Res Rev 21:42–92.

Prochazka A, Gritsenko V, Yakovenko S (2002) Sensory control of locomotion: reflexes versus higher-level control. Adv Exp Med Biol 508:357–367.

Pulver SR, Thirumalai V, Richards KS, Marder E (2003) Dopamine and histamine in the developing stomatogastric system of the lobster *Homarus americanus*. J Comp Neurol 462:400–414.

Roffman RC, Norris BJ, Calabrese RL (2012) Animal-to-animal variability of connection strength in the leech heartbeat central pattern generator. J Neurophysiol 107:1681–1693.

Sachse S, Peele P, Silbering AF, Guhmann M, Galizia CG (2006) Role of histamine as a putative inhibitory transmitter in the honeybee antennal lobe. Front Zool 3:22.

Sauer AE, Buschges A, Stein W (1997) Role of presynaptic inputs to proprioceptive afferents in tuning sensorimotor pathways of an insect joint control network. J Neurobiol 32:359–376.

Schmitz J, Stein W (2000) Convergence of load and movement information onto leg motoneurons in insects. J Neurobiol 42:424–436.

Selverston AI (2010) Invertebrate central pattern generator circuits. Philos Trans R Soc Lond B Biol Sci 365:2329–2345.

Selverston AI, Szucs A, Huerta R, Pinto R, Reyes M (2009) Neural mechanisms underlying the generation of the lobster gastric mill motor pattern. Front Neural Circuits 3:12–12.

Shaw AC, Jackson AW, Holmes T, Thurman S, Davis GR, McClellan AD (2010) Descending brain neurons in larval lamprey: spinal projection patterns and initiation of locomotion. Exp Neurol 224:527–541.

Smarandache CR, Stein W (2005) The influence of a sensory cell on the central pattern generators in the stomatogastric nervous system of the crab. In: Society for Neuroscience Annual Meeting. Washington, D.C.

Smarandache CR, Stein W (2007) Sensory-induced modification of two motor patterns in the crab, *Cancer pagurus*. J Exp Biol 210:2912–2922.

Smarandache CR, Daur N, Hedrich UB, Stein W (2008) Regulation of motor pattern frequency by reversals in proprioceptive feedback. Eur J Neurosci 28:460–474.

Soleng AF, Chiu K, Raastad M (2003) Unmyelinated axons in the rat hippocampus hyperpolarize and activate an H current when spike frequency exceeds 1 Hz. J Physiol 552:459–470.

Städele C, Stein W (2015) The site of spontaneous ectopic spike initiation is regulated to facilitate signal integration in a sensory neuron. J Neurosci under review.

Städele C, Andras P, Stein W (2012) Simultaneous measurement of membrane potential changes in multiple pattern generating neurons using voltage sensitive dye imaging. J Neurosci Methods 203:78–88.

Stein W (2009) Modulation of stomatogastric rhythms. J Comp Physiol A 195:989–1009.

Stein W (2014) Sensory Input to Central Pattern Generators. In: Encyclopedia of Computational Neuroscience (Jaeger D, Jung R, eds), pp 1–11: Springer New York.

Stein W, Schmitz J (1999) Multimodal convergence of presynaptic afferent inhibition in insect proprioceptors. J Neurophysiol 82:512–514.

Stein W, DeLong ND, Wood DE, Nusbaum MP (2007) Divergent co-transmitter actions underlie motor pattern activation by a modulatory projection neuron. Eur J Neurosci 26:1148–1165.

Thiele TR, Donovan JC, Baier H (2014) Descending control of swim posture by a midbrain nucleus in zebrafish. Neuron 83:679–691.

von Philipsborn AC, Liu T, Yu JY, Masser C, Bidaye SS, Dickson BJ (2011) Neuronal Control of *Drosophila* Courtship Song. Neuron 69:509–522.

Wolf H (1991) Sensory feedback in locust flight patterning. In: Locomotor neural mechanism in arthropods and vertebrates. (Armstrong DM, Bush BMH, eds): Manchester University Press.

Wolf H (1995) Plasticity in insect leg motor control: interactions of locomotor program and sensory signal processing. Verh Dtsch Zool Ges 88:153–164.

Wolf H, Burrows M (1995) Proprioceptive sensory neurons of a locust leg receive rhythmic presynpatic inhibition during walking. J Neurosci 15:5623–5636.

Xiong W, Chen WR (2002) Dynamic gating of spike propagation in the mitral cell lateral dendrites. Neuron 34:115–126.

